# ERH as a component of the Microprocessor facilitates the maturation of suboptimal microRNAs

**DOI:** 10.1101/2020.05.13.093278

**Authors:** S. Chul Kwon, Harim Jang, Jihye Yang, Jeesoo Kim, S. Chan Baek, Jong-Seo Kim, V. Narry Kim

## Abstract

The Microprocessor complex cleaves the primary transcript of microRNA (pri-miRNA) to initiate miRNA maturation. Microprocessor is known to consist of RNase III DROSHA and dsRNA-binding DGCR8. Here we identify Enhancer of Rudimentary Homolog (ERH) as a new component of the Microprocessor. ERH binds to a conserved region in the N-terminus of DGCR8. Knockdown of ERH or deletion of the DGCR8 N-terminus results in a decrease of processing of primary miRNAs with suboptimal hairpin structures that reside in polycistronic miRNA clusters. ERH increases the processing of suboptimal pri-miR-451 in a manner dependent on its neighboring pri-miR-144. Thus, the ERH dimer may mediate “cluster assistance” in which the Microprocessor is loaded onto a poor substrate with help from a high-affinity substrate in the same cluster. Our study reveals a role of ERH in the miRNA pathway.

## INTRODUCTION

The canonical maturation pathway of microRNA (miRNA) begins with the transcription of a miRNA gene by RNA polymerase II. The primary transcript (also known as pri-miRNA) is cleaved by an enzymatic complex Microprocessor. After nuclear export, the cleaved hairpin product (called precursor miRNA or pre-miRNA) is further processed by another RNase DICER. The resulting short duplex RNA is then loaded onto the Argonaute (AGO) protein, and one strand of the duplex is discarded, producing the mature miRNA-AGO complex (referred to as RNA-induced silencing complex or RISC) (1).

The Microprocessor is known to consist of two proteins, RNase III DROSHA and dsRNA-binding protein DGCR8 (2-8). DROSHA and DGCR8 are necessary and sufficient to cleave many pri-miRNAs tested in vitro (2,5). Several other proteins have been reported to associate with the Microprocessor and participate in pri-miRNA processing. For example, DEAD-box RNA helicases DDX5 and DDX17 were reported as DROSHA-interacting proteins (5). hnRNPA1 and KSRP were shown to regulate the processing of a subset of pri-miRNAs (9-11). SRSF3 binds to a cis-acting element (CNNC motif) to enhance the efficiency and accuracy of processing (12,13). However, except for DROSHA and DGCR8, none of these proteins are considered as the Microprocessor components because they do not form a stable complex with DROSHA or DGCR8.

A canonical pri-miRNA has characteristic features that are recognized by the Microprocessor, including a ∼35-bp double-stranded stem, an apical loop (>10 bp), and basal single-stranded regions (12,14,15). In addition, four sequence motifs (basal UG, apical UGU, mismatched GHG, and flanking CNNC motifs) enhance the processing efficiency and cleavage specificity (12,13,15,16).

Recent biochemical and structural studies enabled a clear understanding of the processing mechanism. Single-molecular measurement of the Microprocessor complex showed that the complex is a heterotrimer with one molecule of DROSHA and two molecules of DGCR8 (2,8). DROSHA, together with DGCR8, forms an elongated shape and spans the entire hairpin to measure its stem length (∼35-bp) (15,17-19). The RNA-binding affinity mainly comes from DGCR8 rather than DROSHA (2). However, DROSHA constitutes the core of the Microprocessor. The central domain of DROSHA fits into the basal part of pri-miRNA while the double-stranded RNA-binding domain (dsRBD) of DROSHA binds to the mismatched GHG motif to fix the position of pri-miRNA within the Microprocessor (16,18,19). Two RNase III domains form an intramolecular dimer to create the processing center that cleaves pri-miRNA (6).

Here, we sought to identify the interactors of the Microprocessor, and found a small globular protein, Enhancer of Rudimentary Homolog (ERH), as a stable Microprocessor component. Our study reveals that ERH binds to the N-terminus of DGCR8 and enables efficient processing of suboptimal hairpins in polycistronic pri-miRNAs, providing new insights into the pri-miRNA processing model.

## MATERIALS AND METHODS

### Cell lines

HEK293E cells (human embryonic kidney 293 EBNA1; authenticated by ATCC STR profiling) were grown in DMEM (Welgene) supplemented with 9% fetal bovine serum (Welgene). K562 cells (Korean Cell Line Bank; authenticated by ATCC STR profiling) were grown in RPMI1640 (Welgene) supplemented with 9% fetal bovine serum (Welgene).

### Preparation of knock-in cell line by CRISPR/Cas9

CRISPR-Cas9 experiments were designed and conducted based on previous papers (20,21). Briefly, an oligo duplex encoding the sgRNA sequence was cloned at the Bbs I site of pX458-pSpCas9 (BB)-2A-GFP plasmid (Addgene #48138). To construct a donor vector, the ∼600 bp homology sequence to the target site was amplified by overlapping PCR and inserted into a Sal I and Not I digested pGL3-Basic plasmid (Promega). Cas9 and donor plasmids were co-transfected using Lipofectamine 2000 (Thermo Fisher Scientific) into HEK293E cells which were seeded in a 6-well plate one day before transfection. Two days after transfection, cells were diluted to a final concentration of 0.5 cells per 100 μl and plated in two 96-well plates. After 2 weeks, single cell-derived colonies were picked and cultured in a 96-well plate for further experiments. To detect the insertion of the tag sequence by PCR analysis, the genomic DNA of each clone was extracted by using QuickExtract solution (Epicentre) and PCR was carried out. Genomic amplicons of the target region were cloned into a TopBlunt V2 vector (Enzynomics) and verified by Sanger sequencing. Subsequently, the expression of the 3xFlag-2xStrep tag was confirmed by immunoprecipitation and western blotting.

### Small RNA sequencing

Sequencing libraries were generated as previously described (16,22). Briefly, HEK293E cells were tandemly transfected with siRNAs using Lipofectamine 2000 on days 0 and 3. On day 5, total RNA was purified using TRIzol (Thermo Fisher Scientific), and 10 μg of total RNA was used for library preparation. K562 cells were resuspended in OPTI-MEM (Thermo Fisher Scientific) and transfected with siRNAs using Neon (Thermo Fisher Scientific) with the condition of 1450 V, 10 ms, 3 pulses. Four days after transfection, total RNA was purified with TRIzol. Most of the sequencing analysis steps were done as previously described (16). In this paper, all the genome mapped reads (including ambiguously mapped ones) were used for counting miRNA reads. The normalization was done by RPM (reads per million) or DESeq2 (23). Small RNA-seq results from previous papers were downloaded from GEO (GSE141098 (24), GSE116303 (25)) and analyzed through the same pipeline except for handling different adaptors. To filter out lowly expressed miRNAs for reliable analysis, baseMean values from DESeq2 were used for cutoff: 800 (siERH), 100 (DGCR8 Δex2), 50 (SAFB DKO, HEK293T), 100 (SAFB DKO, Ramos). Linear regression models were drawn using seaborn.regplot (size of the confidence interval = 95%).

### Sequence analysis

Protein secondary structure prediction was carried out by using PSIPRED 4.0 (26), and disordered regions were predicted with IsUnstruct v2.02 (27). Conservation scores were calculated by using the Scorecons server (28,29) with the input sequences listed in Supplementary Figure S2. Secondary structures of pri-miRNAs were predicted with RNAfold (30) and then visualized with VARNA (31).

### Immunoprecipitation for mass analysis

Approximately 1.3E9 cells were resuspended in the Lysis buffer (20 mM Tris pH 8.0, 150 mM NaCl, 2 mM beta-mercaptoethanol, 100 ng/μl RNase A) and lysed by sonication. After centrifugation at 35,000 g for 1 hr at 4°C, lysates were incubated with Strep-Tactin beads (IBA Lifesciences) at 4°C overnight. Beads were washed with the Wash buffer (20 mM Tris pH 8.0, 150 mM NaCl, 2 mM beta-mercaptoethanol) for 5 times and incubated with the 50 mM biotin-containing Wash buffer for 1 hr at 4°C. Biotin eluates were further incubated with anti-Flag M2 agarose beads for 3 hr at 4°C. After washing, proteins were eluted with 500 ng/μl of the 3x Flag peptide-containing Wash buffer.

### LC-MS/MS analysis and protein identification

The immuno-precipitated protein samples were first reduced and alkylated in 8 M urea with 50 mM ammonium bicarbonate (ABC) buffer. After 10-fold dilution with 50 mM ABC buffer, the protein samples were digested with 2% (w/w) trypsin at 37 °C overnight. The resulting peptide samples were subject to C18 clean-up and loaded to the in-house packed trap column (3 cm × 150 µm i.d) and capillary analytical column (100 cm x 75 µm i.d.) with 3 µm Jupiter C18 particles (Phenomenex) for peptide separation. A flow rate of 300 nl/min and a linear gradient ranging from 95 % solvent A (water with 0.1% formic acid) to 40% of solvent B (acetonitrile with 0.1% formic acid) for 100 min were applied on nanoACQUITY UPLC (Waters) coupled with Orbitrap Fusion Lumos mass spectrometer (Thermo Fisher Scientific), which was operated at sensitive mode using the following parameters: m/z 350–1800 of precursor scan range, 1.4 Th of precursor isolation window, 30% of normalized collision energy (NCE) for higher-energy collisional dissociation (HCD), 30 s of dynamic exclusion duration, 120k or 30k resolution at m/z 200 and 100 ms or 54 ms of maximum injection time for full MS or MS/MS scan, respectively.

MS raw data files were processed with MaxQuant (version 1.5.3.30) against the human SwissProt database at default settings (20 ppm or 6 ppm of precursor ion mass tolerances for initial or main search, respectively, and 0.5 Da for fragment ion masses). Enzyme specificity was set to trypsin/P and a maximum of two missed cleavages were allowed. Cysteine carbamidomethylation and methionine oxidation were selected as fixed and variable modifications, respectively. A 1% false discovery rate was required at both the protein- and the peptide-level.

### Plasmids

Primers used for cloning are provided in Supplementary Table S3.

### Immunoprecipitation of endogenous proteins

For endogenous protein immunoprecipitation, HEK293E cells grown on 100 mm dishes were lysed in the IP buffer (20 mM Tris pH 7.5, 100 mM KCl, 0.2 mM EDTA, 0.2% NP-40, 10% glycerol, 0.4 ng/μl RNase A and protease inhibitor cocktails (Calbiochem)) and sonicated. Clear lysates were collected by centrifugation for 10 min at 4°C. The antibody was pre-incubated with 20 μl of prewashed protein G Sepharose (GE healthcare) at 4°C for 1 hr. Antibody-conjugated beads were incubated with lysates at 4°C for 1–2 hr and washed with the Wash buffer (20 mM Tris pH 7.5, 100 mM KCl, 0.2 mM EDTA, 0.2% NP-40, 10% glycerol) for 5 times.

### Immunoprecipitation of ectopically expressed proteins

For ectopic expression of the tagged protein (Flag, HA, myc), 10 μg of 1:1 mixture of branched polyethyleneimine (PEI) (Sigma) and linear PEI (Polysciences) were diluted in 400 μl OPTI-MEM with total 5 ug of the expression plasmids. The mixture was incubated for 5 mins before adding to 50% confluent HEK293E cells in a 100 mm dish. After two days, cells were harvested and lysed in the Lysis buffer (20 mM Tris pH 7.5, 150 mM NaCl, 0.1 ng/ul RNase A, and protease inhibitor cocktails (Calbiochem)) by sonication. The cell lysates were collected by full speed centrifugation for 10 min at 4°C.

For Flag or HA-tagged protein immunoprecipitation, anti-Flag M2 affinity gel (Sigma-Aldrich) or anti-HA agarose affinity gel (Sigma-Aldrich) was prewashed three times with 1 ml of the Wash buffer (20 mM Tris pH 7.5, 150 mM NaCl). In the case of anti-myc immunoprecipitation, the antibody was pre-incubated with 20 μl of prewashed protein G sepharose (GE healthcare) at 4°C for 1 hr. Lysates and beads were incubated at 4°C for 1–2 hr. Incubated beads were washed three times with the Wash buffer. Remaining supernatants were removed using a 30 gauge needle. Beads were resuspended in 1X SDS protein sample buffer (50 mM Tris pH 6.8, 10% Glycerol, 2% SDS, 100 mM DTT, 0.01% Bromophenol Blue) for western blotting.

### Western blotting

For western blotting, protein samples were boiled at 95°C for 5 min before separating on Novex WedgeWell 8–16% Tris-Glycine Mini Gels (Thermo Fisher Scientific). Protein samples were transferred to polyvinylidene fluoride membrane (GE Healthcare). The signals were detected using the Odyssey infrared imaging system (Odyssey Sa, LI-COR) or using SuperSignal West Pico PLUS Chemiluminescent Substrate (Thermo Fisher Scientific).

### RNA preparation

Total RNA was prepared using TRIzol (Thermo Fisher Scientific) following the manufacturer’s protocol. To remove remaining DNA contaminants, isolated total RNA was treated with DNase I (Takara) for 0.5–1 hr at 37°C and re-purified using Acid-Phenol/Chloroform (Ambion). In some cases, total RNA was prepared with Maxwell 16 LEV simplyRNA Tissue kit (Promega).

### ERH knockdown and qRT-PCR

To validate the role of ERH in the polycistronic suboptimal miRNA processing, siERH targeting 3′ UTR of ERH was transfected into 50% confluent HEK293E cells using lipofectamine 3000 (Thermo Fisher Scientific) following manufacturer’s instructions. After 48 hours, the cells were trypsinized and counted for reverse transfection. For reverse transfection, 40 pmol of siRNA and 1 μg of pri-miR-144∼451a expression vector were co-transfected into 4E5 cells in a 6-well culture plate using 3 μl of lipofectamine 3000. After additional 48 hours, RNA was harvested with TRIzol.

Relative accumulation of miRNA was calculated by two separate qPCR results. The level of mature miRNAs was measured with the TaqMan MicroRNA Assays (Thermo Fisher Scientifics, #001973 for U6 snRNA, #002676 for miR-144, #001105 for miR-451a) as manufacturer’s instruction. The level of ectopic pri-miRNA transcripts was measured with qRT-PCR using SYBR green MIX (Thermo Fisher Scientific). The cDNAs for the qPCR were synthesized from 500 ng of total RNA using random hexamer primers (Thermo Fisher Scientific) and RevertAid Reverse Transcriptase (Thermo Fisher Scientific). U6 snRNA was used as the internal control for both qPCR. The pri-miRNA expression vector contains an SV40 artificial intron, in which the pri-miRNA sequence is inserted, and the sfGFP coding sequence. Transfection efficiencies between samples were normalized using sfGFP. Primers used in this experiment are provided in Supplementary Table S3.

### ERH rescue experiment

ERH rescue experiment was performed through two repeated transfections. Endogenous ERH was knocked down with shERH targeting 3′ UTR during the first two days. Polycistronic miRNA expression plasmids and shERH plasmids were co-expressed during the next two days.

shERH plasmids were diluted in 500 μl of OPTI-MEM with 15 μl of Fugene HD (Promega) and mixed well with rapid pipetting. The mixture was incubated for 5 min before adding into 50% confluent HEK293E cells. After 48 hours, the cells were treated with trypsin and then collected and counted. Total 2.5 μg of shERH plasmids, pri-miRNA expression plasmids, a wild-type ERH plasmid (1:2:2 ratio) were co-transfected into 4E5 cells in 6-well culture plates using 5 μl of Fugene HD. After 48 hours, the total RNA was harvested.

## RESULTS

### ERH binds to DGCR8

To examine if there is an unknown factor involved in the processing of pri-miRNA, we identified Microprocessor-associated proteins using a recently developed strategy for human protein complexes (20). Briefly, we first knocked-in a segment encoding three consecutive Flag and two Strep tags (3xFlag-2xStrep) downstream of the start codon in the DGCR8 locus using CRISPR-Cas9. The DGCR8 complex expressed at the endogenous level was then isolated by Strep pulldown and biotin elution, followed by Flag immunoprecipitation (IP) and 3xFlag peptide elution (Figure 1A and Supplementary Figures S1A–B). This Strep-Flag tandem affinity purification allowed highly stringent purification of the complex. Only three proteins, DGCR8, DROSHA, and ERH, were specifically detected with high LFQ intensities (Figures 1B– C and Supplementary Table S1). Because we treated RNase A during purification, ERH seems to bind to DGCR8 (or DROSHA) in an RNA-independent manner.

**Figure 1.**
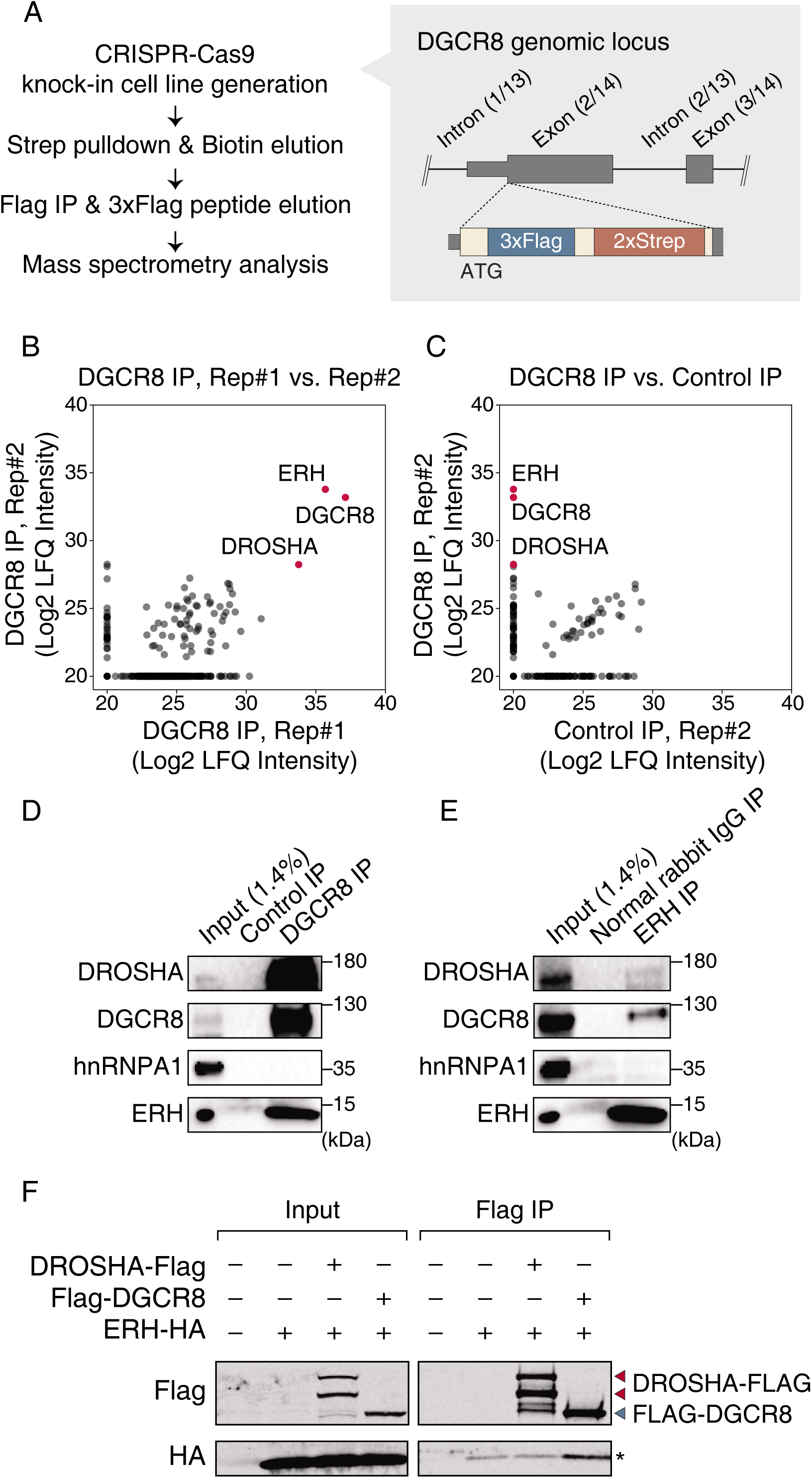
ERH binds to DGCR8. (A) Scheme of 3xFlag-2xStrep-DGCR8 knock-in cell generation, immunoprecipitation, and mass analysis. (B,C) Mass spectrometry results from DGCR8 IP (using the 3xFlag-2xStrep-DGCR8 cell line) and control IP (using the parental cell line). (D,E) Western blots showing the interaction between endogenous DGCR8 and ERH using wild-type HEK293E. The ascites fluid from the SP2/0 fusion partner cell was used for Control IP in D. (F) Western blots with overexpressed DROSHA or DGCR8. Because ectopic DROSHA is aggregated in the absence of C-terminal tail (CTT) of DGCR8 (2,17), DROSHA-Flag was co-transfected with DGCR8 CTT. Some overexpressed ERH proteins were nonspecifically pulled down by anti-Flag agarose beads and denoted with an asterisk (*).

ERH was initially identified as a recessive enhancer of a weak wing phenotype of the rudimentary mutant in *Drosophila melanogaster* (32). Diverse functions of ERH have been described in many organisms. In *Xenopus*, ERH was reported as a cell type-specific transcriptional repressor (33) while in *Schizosaccharomyces pombe*, Erh1 promotes meiotic mRNA decay during vegetative growth (34). Human ERH was shown to be involved in chromosome segregation during mitosis (35,36).

The structure of ERH may provide an insight into how a conserved ERH protein family can perform multiple functions. ERH is a small globular protein (∼12 kDa) that forms a homodimer (37-39). In *S. pombe*, a dimerized Erh1 binds and connects two molecules of Mmi1, a YTH domain-containing RNA-binding protein, and the dimerization is required for the in vivo activity of Mmi1 (40,41). Thus, the functions of ERH may derive from the modulation of each binding partner.

We validated the interaction between DGCR8 and ERH by co-IP (in the presence of RNase A) and western blotting. First, endogenous DGCR8 was pulled down with a mouse monoclonal antibody, and we could detect endogenous ERH from the precipitant (Figure 1D). Conversely, the endogenous ERH complex was immuno-purified, and endogenous DGCR8 was co-precipitated (Figure 1E). In this condition, hnRNPA1, a highly expressed nuclear RNA-binding protein, was not precipitated with either DGCR8 or ERH (Figures 1D and E). Also, it is noted that endogenous DROSHA was also detected in the ERH immuno-purified sample (Figure 1E), implying that ERH binds to Microprocessor.

To determine whether ERH binds to DGCR8 or DROSHA, we performed co-IP using ectopically expressed HA-tagged ERH and Flag-tagged DGCR8 or DROSHA. Since overexpressed DROSHA aggregates in the absence of the partner protein DGCR8, we co-transfected the C-terminal tail (CTT) of DGCR8 as previously described (2). CTT is a short alpha-helix (23 aa) that binds and stabilizes DROSHA (17,42). Notably, ERH was enriched in the Flag-DGCR8 precipitates, but not in the DROSHA-Flag precipitates (Figure 1F), suggesting that ERH binds to DGCR8 rather than DROSHA.

### ERH binds to the N-terminus of DGCR8

The molecular function of each domain of DGCR8 has been extensively studied. RNA-binding heme domain (RHED) mediates homodimerization and recognizes the apical loop of pri-miRNA (43,44). Two double-stranded RNA-binding domains (dsRBDs) bind to the upper stem of pri-miRNA and are responsible for the major RNA-binding affinity of Microprocessor (2,19). However, the function of the N-terminal region of DGCR8 is still unknown. The N-terminus of DGCR8 contains regions that are conserved across vertebrates, suggesting an important contribution (Figure 2A and Supplementary Figure S2A).

**Figure 2.**
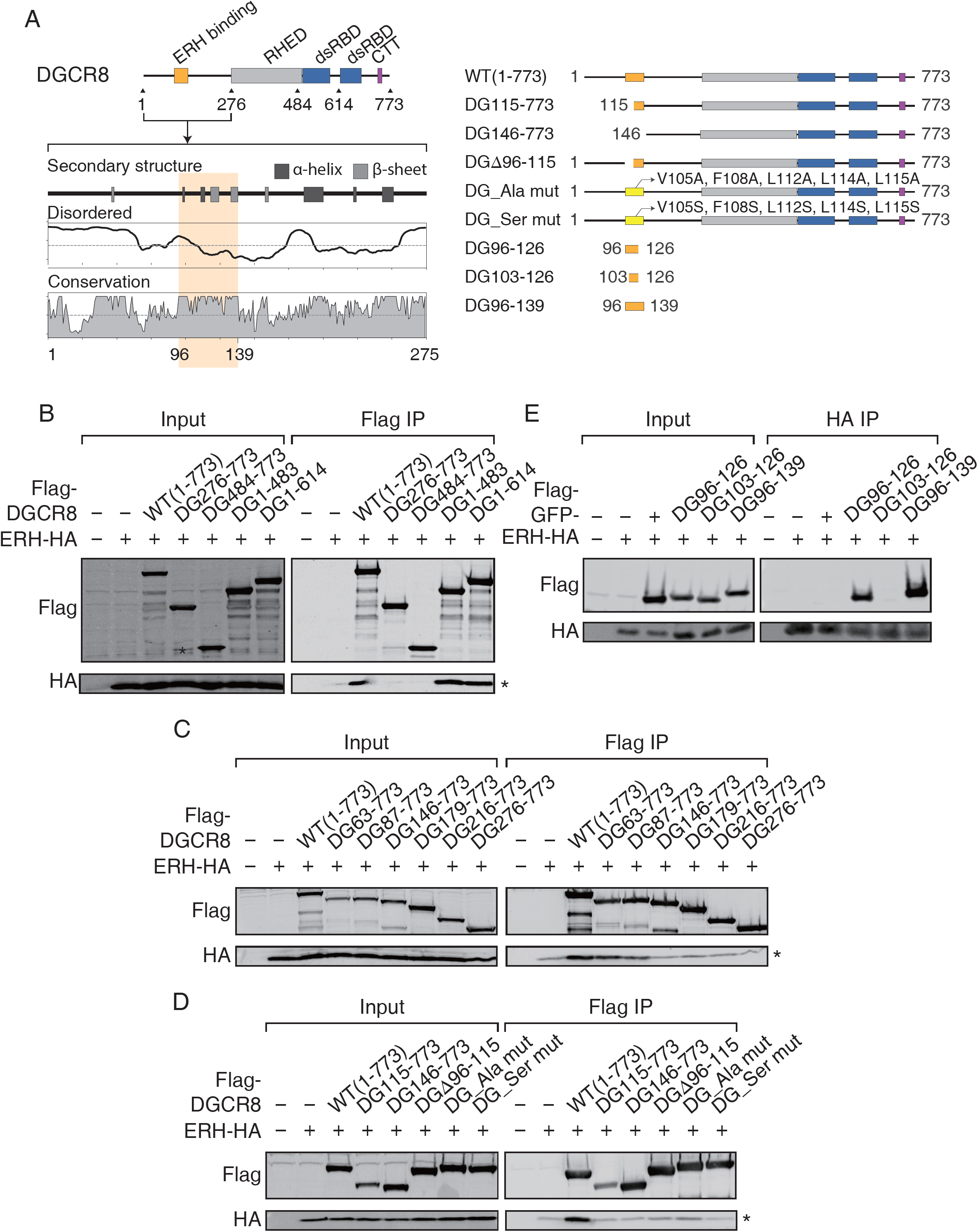
ERH binds to the N-terminus of DGCR8. (A) Sequence analysis of the N-terminus of human DGCR8. (B–D) Western blots with deletion or point mutants of DGCR8. Some overexpressed ERH proteins were nonspecifically pulled down by anti-Flag agarose beads and denoted with an asterisk (*). (E) Western blots with GFP-fused DGCR8 fragments.

To find the ERH-binding site of DGCR8, we tested the interaction between ERH and serial deletion mutants of DGCR8. The deletion mutants that contain the N-terminus of DGCR8 (1–275 aa) pulled down the ERH protein (Figure 2B). We narrowed down the interface between DGCR8 and ERH using additional N-terminal deletion mutants and found that the region between Ile87 and Ala146 is required for binding to ERH (Figure 2C).

The region between Pro96 and Gly139 is highly conserved among vertebrates (Figure 2A, highlighted in yellow and Supplementary Figure S2A). We further prepared several DGCR8 mutants that do not disrupt the overall protein folding based on the predicted secondary structure. Initially, we focused on the region between Pro96 and Leu115 because there were some conserved hydrophobic residues that might be used for protein docking (Supplementary Figure S2A). We deleted the region by generating DG115–773 (where 1–114 aa of DGCR8 was deleted) and DGΔ96–115 (where 96–115 aa of DGCR8 was deleted). These two mutants could not pull down ERH, confirming the importance of this region in the interaction with ERH (Figure 2D). Moreover, we tested additional mutants in which five hydrophobic residues within the 96–115 aa of DGCR8 were replaced with alanine or serine (DG_Ala mut: V105A/F108A/L112A/L114A/L115A; DG_Ser mut: V105S/F105S/L112S/L114S/L115S). These two DGCR8 mutants were impaired in ERH binding, indicating that the hydrophobic residues participate in ERH binding (Figure 2D). Single-point mutations in each hydrophobic residue had little effect on the affinity to the ERH protein compared to that of wild-type (Supplementary Figure S2B), suggesting that the binding interface is formed via multiple weak interactions.

To identify a necessary and sufficient binding site of DGCR8 against ERH, we prepared three short DGCR8 fragments fused to the C terminus of GFP (Figure 2E). Interestingly, DG103–126 that contains the five hydrophobic residues we tested, did not bind to ERH, but DG96–126 was able to interact with ERH (Figure 2E). Further, DG96–139 binds more avidly to the ERH protein compared to DG96–126 (Figure 2E), indicating that the whole conserved block of DGCR8 (Pro96–Gly139) serves as the ERH-binding domain. The hydrophobic residues, as well as additional neighboring residues, are necessary for the binding between DGCR8 and ERH.

### Suboptimal pri-miRNAs are regulated by ERH and the N-terminus of DGCR8

We next investigated whether ERH affects the biogenesis of miRNAs. We performed small RNA-seq after ERH knockdown in HEK293E and K562 cells. Some miRNAs were significantly altered (Figures 3A–C). For validation, we carried out qRT-PCR on pri-miRNAs and compared the results with small RNA-seq. Upon depletion of ERH, the levels of pri-miR-196a-1 and pri-miR-197 modestly decreased while those of mature miR-196a-1 and miR-197 increased (Figure 3D). On the contrary, pri-miR-181b-1 and pri-miR-425 were markedly up-regulated while their mature counterparts were down-regulated strongly in ERH depleted cells (Figure 3D). These data suggest that ERH plays a role in the pri-miRNA processing of a subset of miRNAs.

**Figure 3.**
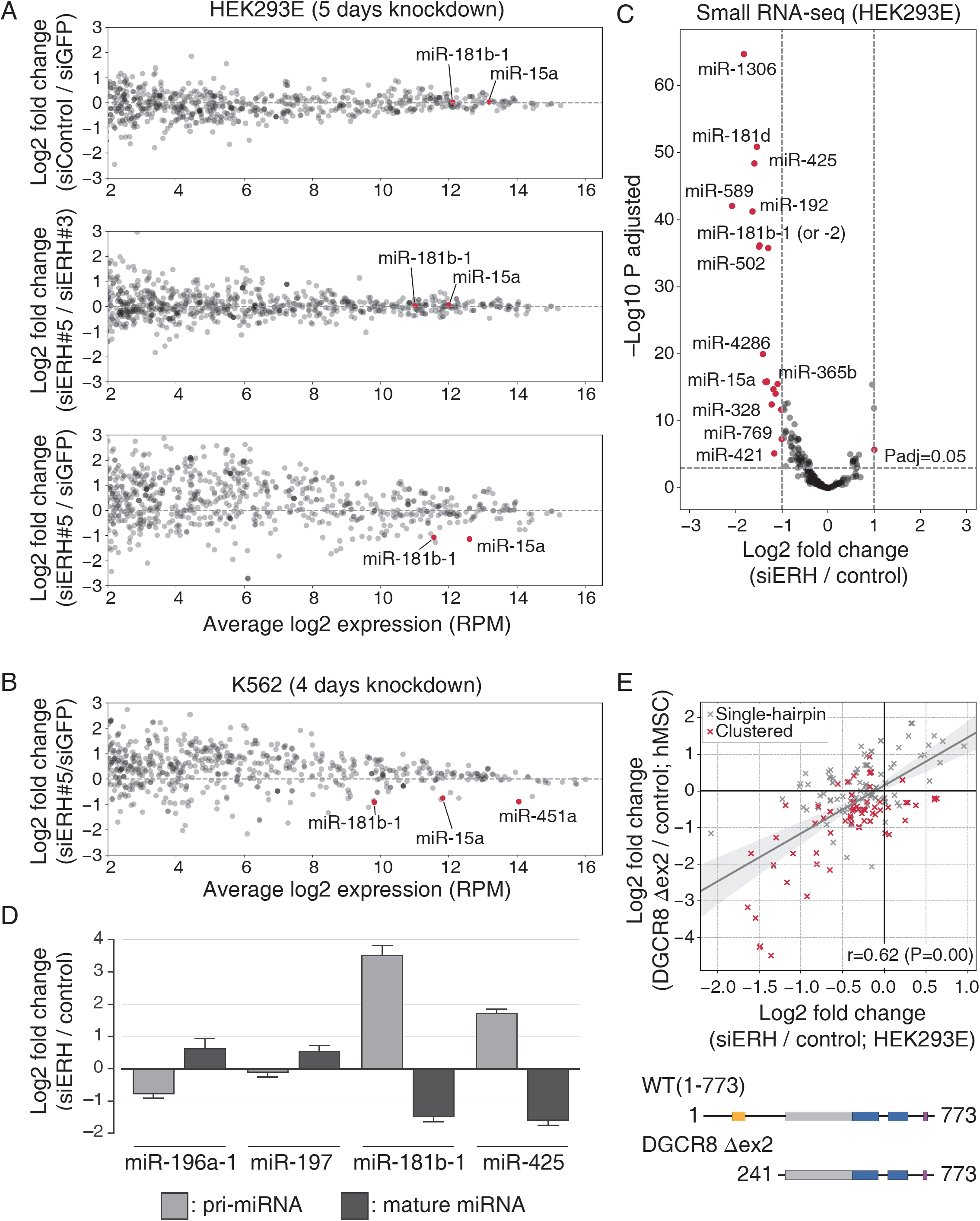
Suboptimal pri-miRNAs are regulated by ERH and the N-terminus of DGCR8. (A,B) MA plots showing the expression of miRNAs mapped on each pri-miRNA genomic locus from ERH-depleted HEK293E cells (A) or K562 cells (B). Representative miRNAs are shown in red. hsa-mir-451a is not significantly expressed in HEK293E. (C) Volcano plot showing differentially expressed miRNAs from ERH-depleted HEK293E cells. (D) qRT-PCR for validation of small RNA-seq. The expression levels of pri-miRNAs were detected by SYBR Green-based qRT-PCR after knocking down ERH for 3 days, and those of mature miRNAs were deduced from small RNA-seq. Error bars indicate standard error of the mean (n = 3) (E) Scatter plots of log2 fold changes of siERH (this study) vs. DGCR8 Δex2 (25). A miRNA is defined as a “clustered” one if another miRNA gene is found within ±1,500 bp in the genome. r is Pearson’s correlation coefficient.

Recently, the Guang-Hui Liu group generated a cell line with a deletion of the DGCR8 N-terminal 240 aa (DGCR8 Δex2; deletion of exon 2) in human mesenchymal stem cells (25). The deletion resulted in differential expression of certain miRNAs as determined by small RNA-seq (25). We noticed that the deleted region of DGCR8 (1–240 aa) encompasses the ERH-binding site (96–139 aa), and therefore, we hypothesized that the miRNA expression profiles might be consistent. Indeed, even though the cell lines are different, there was a correlation between commonly expressed miRNAs (Figure 3E). Intriguingly, many commonly down-regulated miRNAs are polycistronic pri-miRNAs (Figure 3E, colored in red). Taken together, these data implicate that ERH and the N-terminus of DGCR8 may act together in the processing of polycistronic pri-miRNAs.

We further examined the features of miRNAs whose expression was strongly impaired upon ERH knockdown. Notably, these miRNAs have unusual hairpin structures that are unlikely to be efficiently processed by Microprocessor (Supplementary Figure S3). For example, miR-1306 is produced from a hairpin that resides in the second exon of DGCR8 (45,46). The main function of this hairpin is to destabilize the DGCR8 mRNA so as to achieve an autoregulatory homeostatic control of Microprocessor activity. So the miR-1306 hairpin is an inherently suboptimal substrate that can be cleaved only when the Microprocessor level is high (45,46). pri-miR-15a and pri-miR-181b-1 have a large unpaired lower stem structure while the basal flanking segments of pri-miR-425 are partially double-stranded (Supplementary Figure S3). pri-miR-192, pri-miR-328, and pri-miR-589 also seem to have suboptimal structures in the lower stem and the apical loop (Supplementary Figure S3).

pri-miR-451a, an extensively studied suboptimal hairpin (47,48), was also identified as an ERH-affected miRNA in K562 cells (Figure 3B). Many of the pri-miRNAs mentioned above, including miR-451a, have neighboring pri-miRNA hairpins in the same transcripts, suggesting that ERH may affect the processing of suboptimal pri-miRNAs in polycistronic transcripts.

About one-third of human miRNAs reside in “polycistronic clusters” with more than one hairpins in a single nascent transcript (49,50). Intriguingly, the processing of some hairpins depends on that of the neighboring hairpins in the same transcript (51,52), indicating that there is an interplay between clustered hairpins. In a recent study, the Lai group and our laboratory revealed that the processing of pri-miR-451a, a suboptimal substrate for Microprocessor, is enhanced by the neighboring optimal hairpin, pri-miR-144 (47). The Bartel group also independently made a similar observation and showed that ERH facilitates cluster assistance (48). Moreover, before these two papers were published, the Herzog group reported that pri-miR-15a could be processed by the help of the neighbor pri-miR-16-1 and a trans-acting protein SAFB2 (24).

### ERH is required for the cluster assistance of pri-miR-144∼451a

To experimentally verify the role of ERH in the processing of clustered suboptimal pri-miRNAs, we chose the pri-miR-144∼451a cluster (47,48). We generated a construct containing both hairpins or those with a single hairpin (Figures 4A and B). siERH or control siRNA was transfected along with the constructs into HEK293E cells where endogenous pri-miR-144∼451a is rarely expressed. As expected from earlier findings (47,48), a higher level of mature miR-451a is expressed from the miR-144∼451a constructs than from the stand-alone miR-451a construct (Figure 4B). Importantly, in ERH-depleted cells, the mature miR-451a level from the miR-144∼451a construct was drastically reduced (Figure 4B). Moreover, when ERH is ectopically expressed in ERH-depleted cells, it effectively increases the expression of miR-451a without significantly altering miR-144 expression (Figure 4C). Taken together, our data demonstrates a role of ERH in cluster assistance of suboptimal pri-miRNAs.

**Figure 4.**
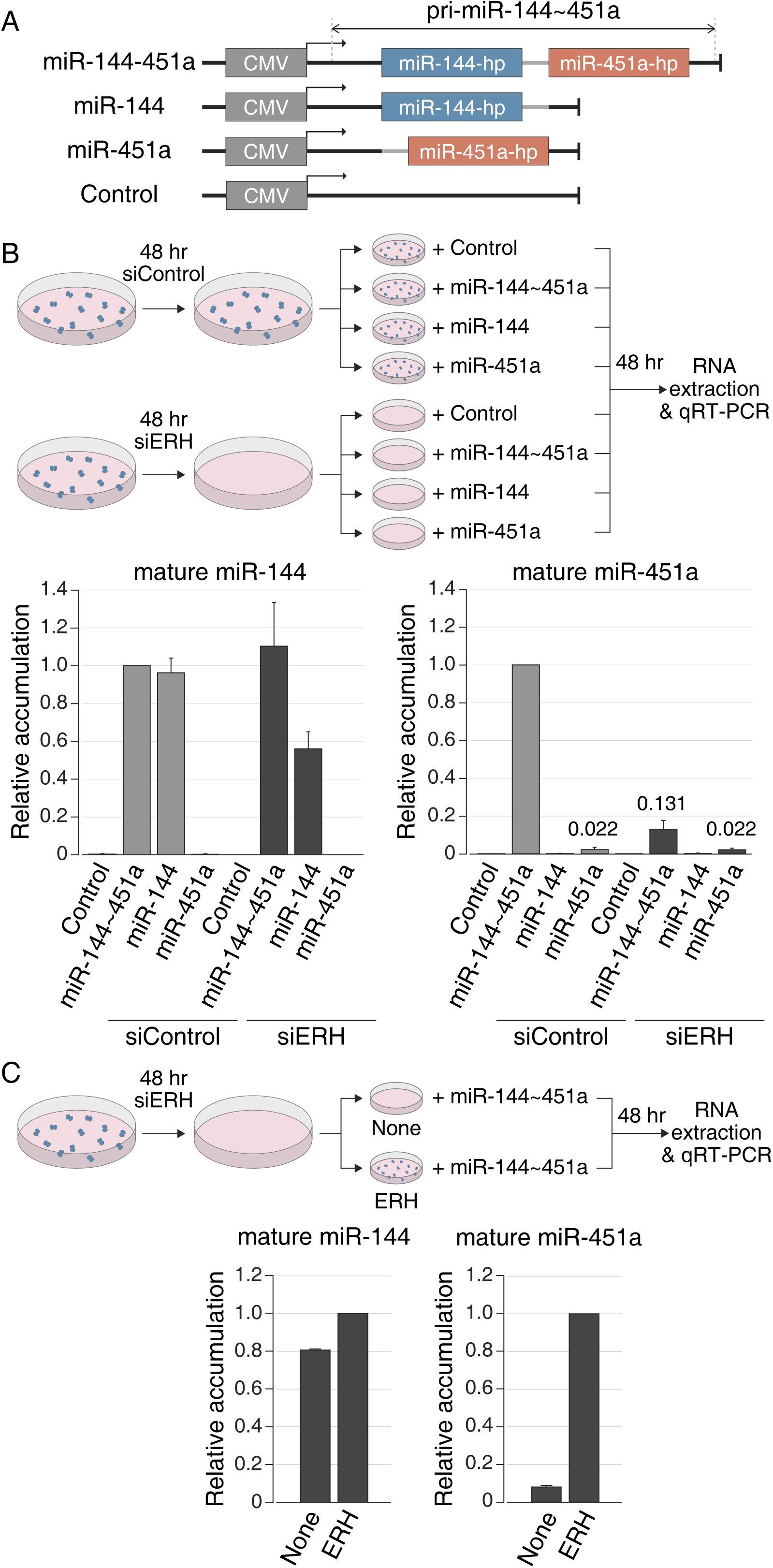
ERH is required for the processing of suboptimal clustered pri-miR-451a. (A) Scheme of ectopic pri-miR-144∼451a constructs. (B,C) qRT-PCR showing the ERH dependency of pri-miR-144∼451a processing. Error bars indicate standard error of the mean (n = 3).

## DISCUSSION

This study introduces ERH as the third core component of Microprocessor. Stringent tandem affinity purification of DGCR8 from the endogenous genomic context yielded only three proteins: DGCR8 (the bait), DROSHA (the well-known partner of DGCR8), and ERH (Figures 1B and C). From literature search, we noticed that the interaction between DGCR8 and ERH has been detected in several previous proteomic experiments even though the results were not validated (53,54). Moreover, the theoretical molecular weight of the complex is close to the experimentally measured weight of the Microprocessor when ERH is considered. Previously, we estimated the mass of an endogenous Microprocessor as ∼364 (±25) kDa using a sedimentation coefficient and a Stokes radius (2). The theoretical weight of the complex is 355 kDa if the complex consists of one molecule of DROSHA (159 kDa), two molecules of DGCR8 (86+86 kDa), and a dimer of ERH (12+12 kDa). Thus, the Microprocessor may be a heteropentamer. Since several proteins besides DGCR8 have been shown to interact with ERH in human cells (54), we propose that there are various ERH-containing complexes, and the Microprocessor is one of them.

We note that while we were still investigating the action mechanism of ERH, the Bartel group reported that ERH is co-purified with the Microprocessor and is involved in cluster assistance (48). In our study, we not only independently found ERH as a co-purified protein but further revealed that ERH directly binds to the N-terminus (96–139 aa) of DGCR8, among which five amino acids (Val105, Phe108, Leu112, Leu114, and Leu115) are critical for the interaction.

Recently, the Herzog group identified SAFB2 as a factor involved in cluster assistance through genome-wide CRISPR screening (24). Interestingly, among the unvalidated positive hits, we could find ERH, implying that both ERH and SAFB2 are required for cluster assistance. To test this possibility, we compared the miRNA alteration patterns of siERH-treated HEK293E cells (this study), the SAFB/SAFB2 double knock out (DKO) Ramos or HEK293T cells (24), and the DGCR8 Δex2 hMSCs (25). The overall alteration patterns were notably similar, and many of the commonly down-regulated pri-miRNAs reside in clusters (Figure 5 and Supplementary Figure S4). These data suggest that ERH, SAFB2, and the N-terminus of DGCR8 play a role together in the cluster-assisted processing of suboptimal hairpins.

**Figure 5.**
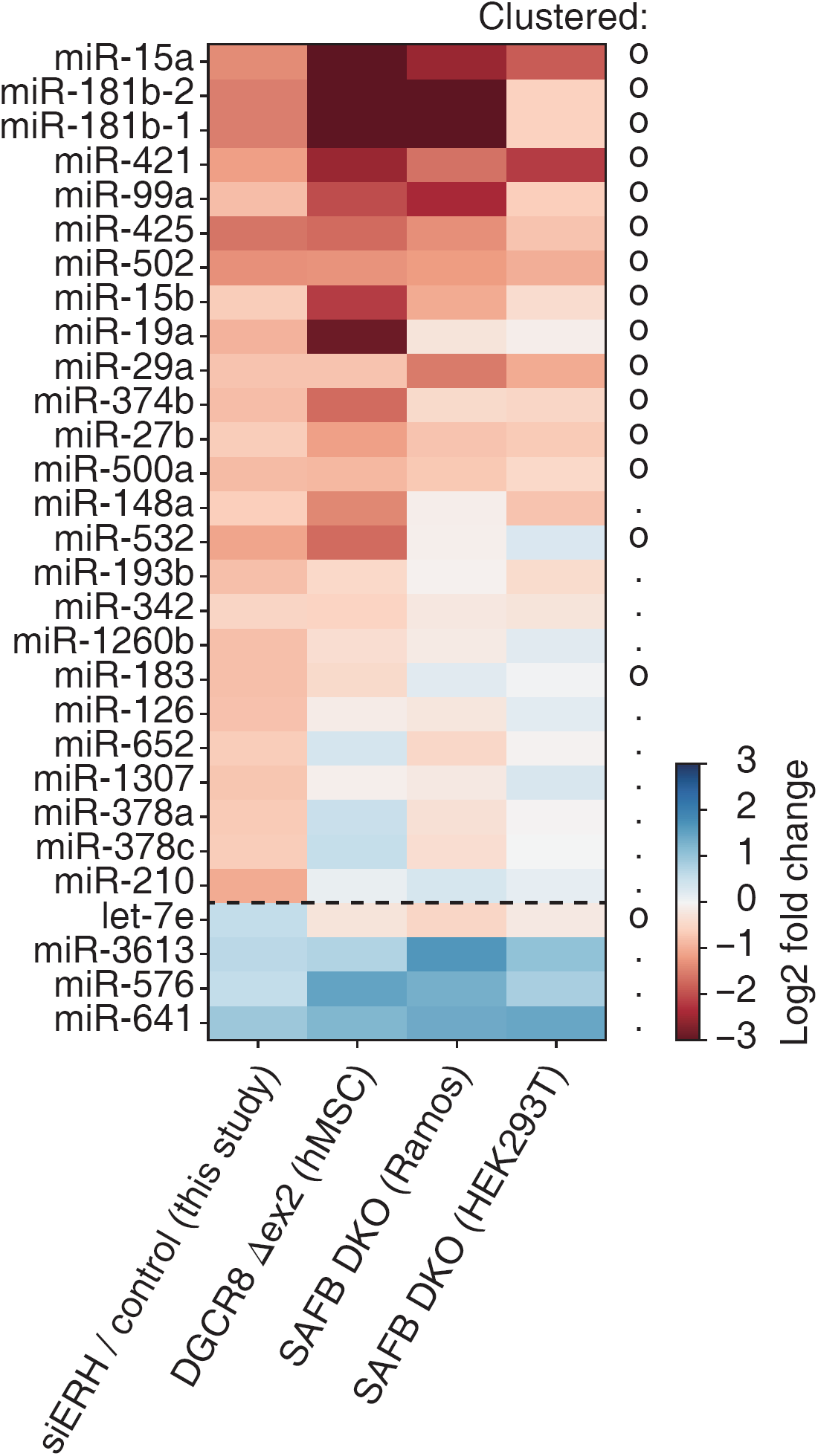
ERH, SAFB2, and the N-terminus of DGCR8 play a role together in the cluster-assisted processing of suboptimal hairpins. Heatmap for comparing small RNA-seq results from siERH, DGCR8 exon 2 deletion, and SAFB/SAFB2 double knock out (DKO). Commonly expressed miRNAs in all samples were first selected, and then differentially expressed miRNAs in the siERH sample (log2 fold change >0.5 or <–0.5) were shown.

Other candidate genes from the CRISPR screening (24) provide additional insight into the mechanism of cluster assistance. First, there were positive hits on miR-1306 hairpin which resides in the second exon of the DGCR8 gene. Since Cas9 induces a short deletion to a target gene, it is likely that the DGCR8 protein was still expressed with a small deletion, and that ERH cannot bind to this DGCR8 as in the DGCR8 Δex2 (Figure 3E). Second, the screening also found UROD (also known as uroporphyrinogen decarboxylase), which is an essential enzyme involved in the heme biosynthetic process. Because heme is an essential cofactor for the dimerization of RHED domain of DGCR8 and the processing of pri-miRNA(43,44), it is likely that the defect of UROD is related to the dimerization of DGCR8. Combining these results, we propose two possible mechanistic models to explain the function of ERH and the mechanism of cluster assistance (Figure 6).

**Figure 6.**
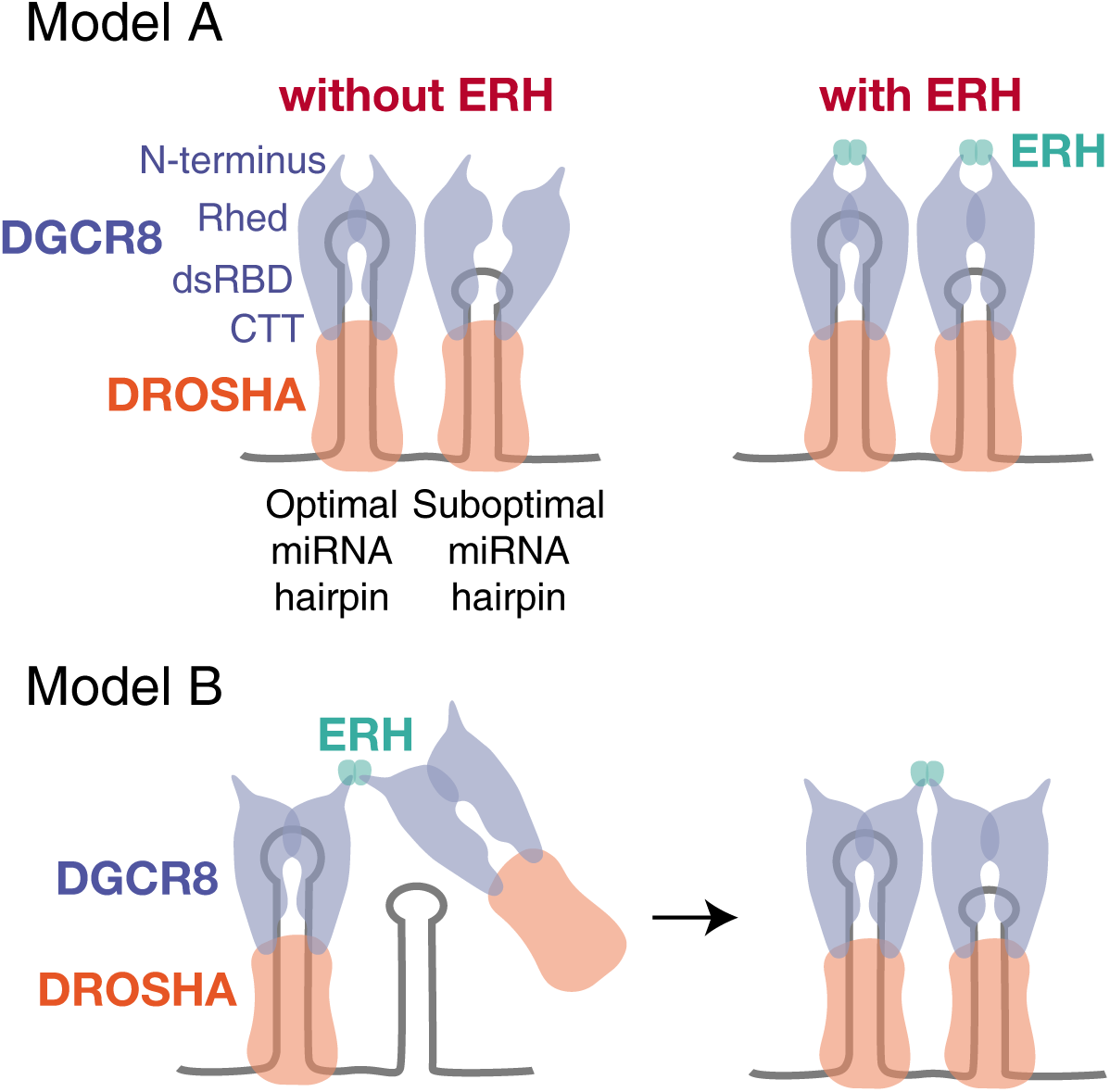
Models of the ERH-mediated cluster-assisted processing of suboptimal pri-miRNAs.

In model A, ERH modulates the interaction between two DGCR8 subunits in the Microprocessor (“intra-complex dimerization of DGCR8 by ERH”) (Figure 6). In this model, the dimerization of the Microprocessor is mainly enabled by SAFB2, which can bind to the N-terminus of DROSHA and may be self-dimerized through the coiled-coil domain (24). Two connected Microprocessors recognize two pri-miRNA hairpins in vicinity. The suboptimal hairpin may not be recognized by an ERH-depleted Microprocessor because two DGCR8 molecules are not accurately aligned.

In contrast, in model B, the ERH dimer connects two Microprocessors (“inter-complex dimerization of DGCR8 via ERH”) (Figure 6). Because the interaction between ERH and SAFB have been previously reported (55), we may also need to consider a slightly modified model where the large SAFB2 protein encompasses the whole Microprocessor units by interacting with both DROSHA and ERH. Two models may not be mutually exclusive, and both models may work in cells. The detailed mechanism of DROSHA, DGCR8, ERH, and SAFB2 needs further investigation using in vitro reconstitution with recombinant proteins.

Some miRNAs that were affected by ERH knockdown do not seem to have neighboring miRNA hairpins. For example, miR-589 and miR-328 have the nearest miRNAs at 215,923 bp and 1,031,031 bp away, respectively (Figure 3C and Supplementary Table S2). Therefore, it is yet to be examined if there is an unknown DROSHA-recruiting RNA element near these miRNA loci.

ERH is a highly conserved protein identified in most eukaryotes and modulates the function of partner proteins (56). However, the ERH-binding site of DGCR8 is only present in vertebrates. Thus, it is likely that ERH has recently joined the miRNA biogenesis pathway and contributed to the emergence of new miRNAs in complex organisms through cluster assistance. Our study provides an example of how the conserved protein ERH plays a unique role in a specific phylogenetic lineage.

## DATA AVAILABILITY

The small RNA-seq data in this study are available at the NCBI Gene Expression Omnibus (accession number: GSE149421).

## SUPPLEMENTARY DATA

Supplementary Data are available at NAR Online.

## ACKNOWLEDGEMENTS

We thank Eunji Kim and Daeun Choi for the technical help, and Young-suk Lee for the critical reading of the manuscript.

## AUTHORS CONTRIBUTIONS

S.C.K. and H.J. performed biochemical experiments. S.C.K. and S.C.B. carried out bioinformatics. J.Y. generated knock-in cell lines. J.K. and J.-S.K. performed mass spectrometry experiments and analysis. S.C.K., H.J. and V.N.K. analyzed the data and wrote the manuscript.

