## Supplementary Figures for "ERH as a component of the Microprocessor facilitates the maturation of suboptimal microRNAs"

### Figure S1

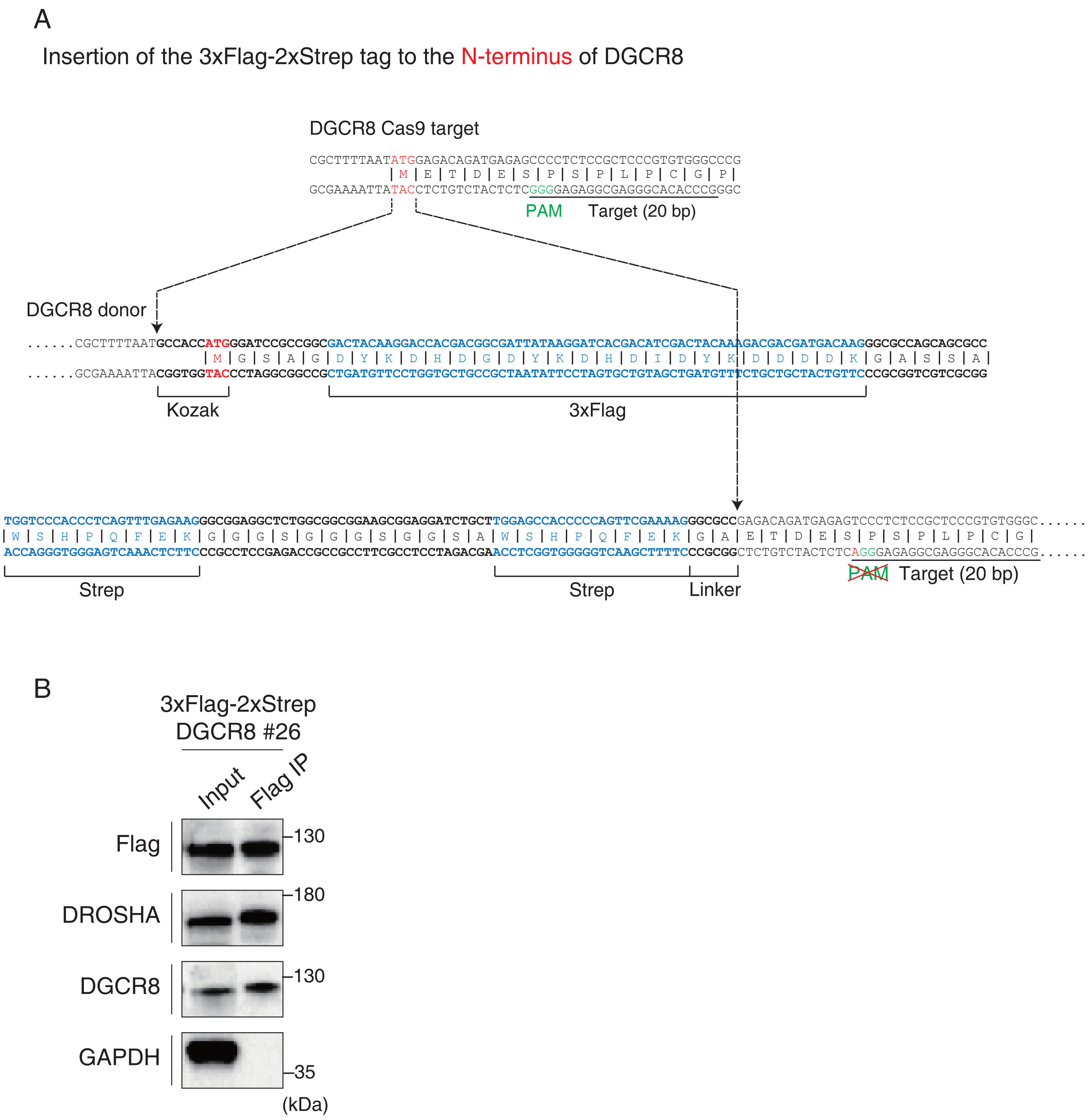


Supplemental Figure S1. (A) Scheme of the DGCR8 locus and donor sequence. (B) Western blots showing the expression of 3xFlag-2xStrep-DGCR8 from the knock-in cell line #26.

Figure S2

A

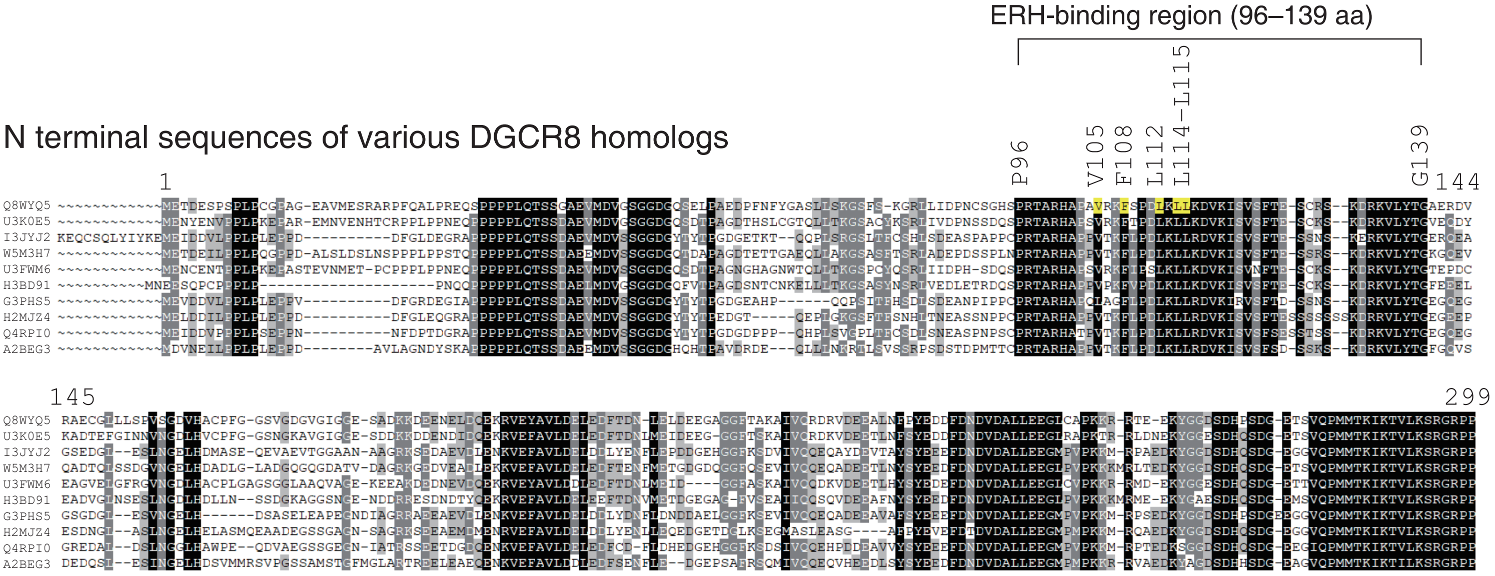


Q8WYQ5 Human DGCR8
U3K0E5 Collared flycatcher
I3JYJ2 Nile tilapia
W5M3H7 Spotted gar
U3FWM6 Eastern coral snake
H3BD91 Coelacanth
G3PHS5 Three-spined stickleback
H2MJZ4 Japanese rice fish
Q4RPI0 Spotted green pufferfish
A2BEG3 Zebrafish

B

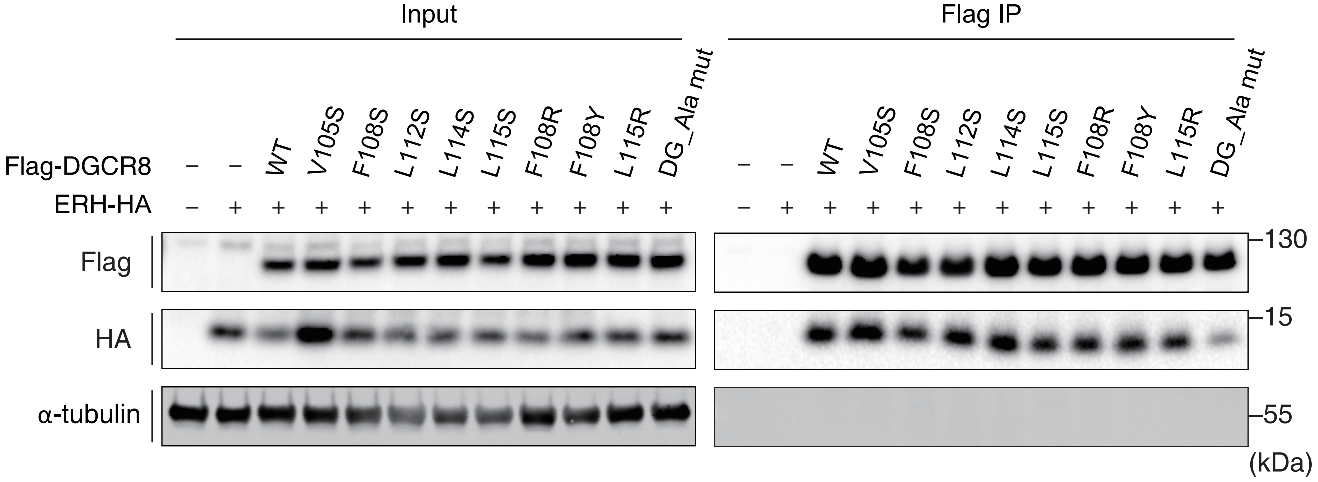


Supplemental Figure S2. (A) Multiple sequence alignment of the N terminus of DGCR8 orthologs. (B) Western blots showing the interaction between ERH-HA and Flag-DGCR8 containing single-point mutation on the ERH-binding site. DG_Ala mut: V105A/F108A/L112A/L114A/L115A.

Figure S3

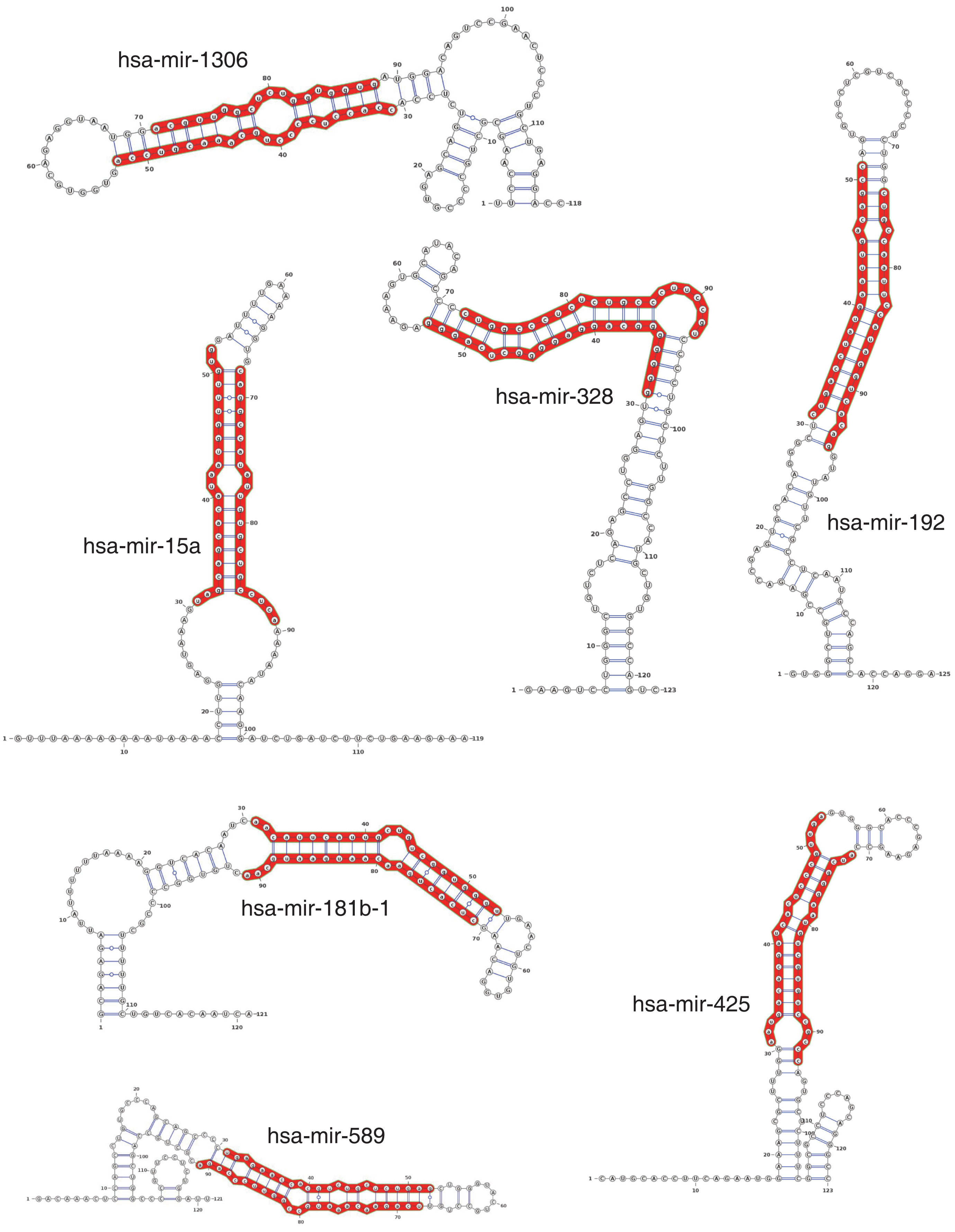


Supplemental Figure S3. The predicted secondary structures of pri-miRNAs affected by siERH. Mature miRNA sequences annotated in miRBase are shown in red.

### Figure S4

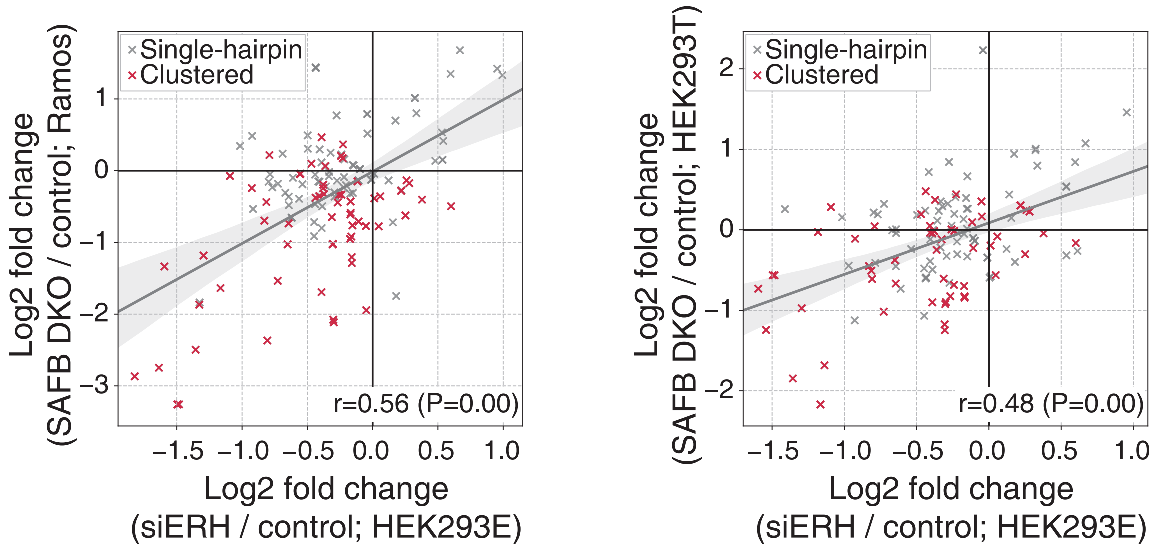


Supplemental Figure S4. Scatter plots of log2 fold changes of siERH (this study) vs. SAFB/SAFB2 double knock out (DKO) (Hutter et al., 2019). A miRNA is defined as a “clustered” one if another miRNA gene is found within ±1,500 bp in the genome. r is Pearson’s correlation coefficient.
